# V_0_-ATPase downregulation induces MVID-like brush border defects independently of apical trafficking in the mammalian intestine

**DOI:** 10.1101/2022.11.04.515188

**Authors:** Aurélien Bidaud-Meynard, Ophélie Nicolle, Anne Bourdais, Maela Duclos, Jad Saleh, Frank Ruemmele, Henner F Farin, Delphine Delacour, Despina Moshous, Grégoire Michaux

## Abstract

**Summary:** Intestinal microvillus atrophy is a major cause of enteropathies such as idiopathic or congenital diarrhea that are often associated with severe morbidity. It can be caused by genetic disorders, inflammatory diseases, toxins or pathogens. In particular, Microvillus inclusion disease (MVID) is characterized by a chronic intractable diarrhea and a severe microvillus atrophy. It is triggered by mutations in *MYO5B, STX3, MUNC18*.*2* or *UNC45A* which alter epithelial polarity by affecting apical trafficking in intestinal epithelial cells. Furthermore, we recently established that the depletion of the V_0_ sector of the V-ATPase complex induces an MVID-like phenotype in *C. elegans*. In this study we investigated the function of the V_0_-ATPase complex in mouse intestinal organoids. We found that its depletion also triggers a very severe microvillus atrophy in this model. Furthermore, we established that the polarity of intestinal cells is affected in a patient carrying mutations in *TCIRG1* which encodes a V_0_-ATPase subunit. However, V_0_- ATPase depletion does not recapitulate other MVID-specific phenotypes such as subapical vesicle accumulation and Rab11+ endosomes mislocalization. Finally, we found that the apical localization of the V_0_-ATPase is disrupted in MVID patients. Altogether these results suggest a role for the V_0_-ATPase in microvillus atrophy which might be independent from apical trafficking.

---

Microvillus inclusion disease (MVID, OMIM 251850) is a rare genetic orphan condition associated with a chronic intractable diarrhea and nutrient absorption defects that compromise the survival of new-born^1^. Mutations found in *MYO5B, STX3, STXBP2*/*MUNC18*.*2* or *UNC45A* in MVID patients^1, 2^ highlighted the role of apical trafficking of transporters and ion channels^3^ in the absorptive function. It also suggested that apical trafficking is involved in the maintenance of the enterocyte brush border (BB), whose atrophy is a typical phenotype of MVID.

We recently demonstrated that knockdown of subunits of the V_0_ sector of the V-ATPase complex (aka V_0_-ATPase) induces an MVID-like phenotype in *C. elegans*^4^. Here, to study the function of V-ATPase in mammals, we depleted V_0_ (*Atp6v0d1, Atp6v0c*) and V_1_ (*Atp6v1e2*) subunits by inducible CRISPR-CAS9 knockout (KO) in mouse intestinal organoids^5^, and analyzed the resulting phenotypes that we compared to that of *Myo5b* depletion, a *bona fide* MVID model^6^.

*Atp6v0d1* but not *Atp6v1e2* KO induced a very severe BB atrophy with smaller, sparse, and slightly wider microvilli as observed by transmission electron microscopy (TEM), similarly to *Myo5b* KO (Figures 1A-B and Supplementary Figure 1A-C). F-Actin staining also revealed the accumulation of cytoplasmic actin^+^ foci in both *Myo5b* and *Atp6v0d1* KO organoids, reminiscent of microvillus inclusions (MVIs)^7^, a typical phenotype of MVID^1^ (Figure 1C). Super-resolution and TEM imaging indeed revealed the presence of MVIs lined with microvilli in *Myo5b, Atp6v0d1* as well as in *Atp6v0c* KO organoids (Figure 1D and Supplementary Figure 1D-E). Ultrastructural analysis showed that both *Atp6v0d1* and *Myo5b* KO induced other MVID phenotypes, such as the accumulation of large vacuoles with heterogeneous content (“mixed- organelles”) and the formation of ectopic lumen between lateral membranes, indicating epithelial polarity defects (Figure 1E and Supplementary Figure 1F-G). Other MVID features such as defective lysosomes and digitations at the basolateral membrane were also observed upon depletion of *Atp6v1e2*, which only inhibits the acidification but not the trafficking function of the V-ATPase^4, 8^. These findings indicate that some MVID hallmarks could be linked to a general defect in organelle pH or autophagy, as suggested before^1^ (Supplementary Figure 1A,H and Supplementary Table1).

**Figure 1.**
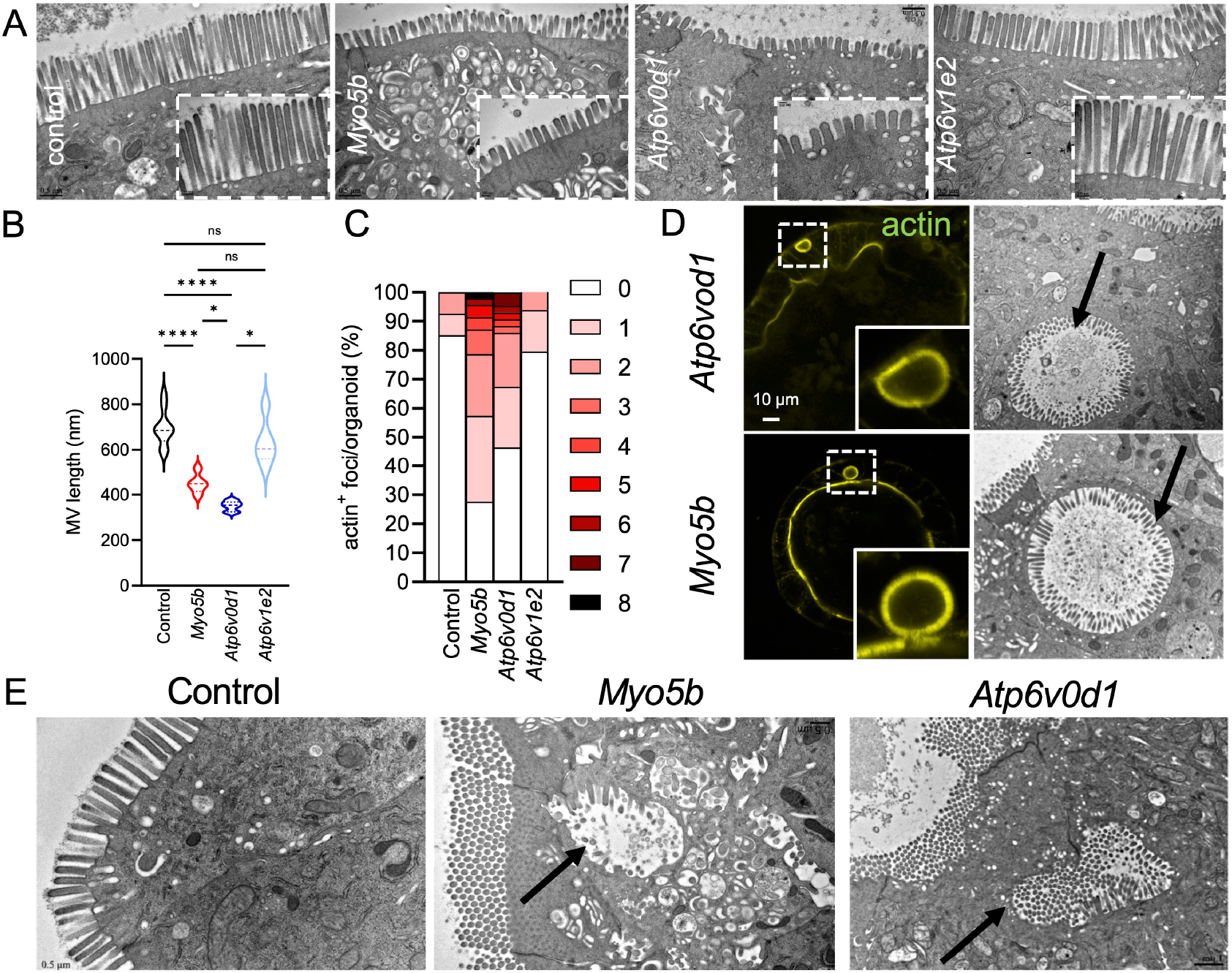
V_0_-ATPase KO recapitulates the MVID-related structural defects. Control organoids and organoids KO for *Myo5b or* the V-ATPase subunits *Atp6v0d1* and *Atp6v1e2* were analyzed by TEM (A-B, D-E) or super-resolution microscopy of F-actin staining (C-D). (B) Quantification of the microvilli (MV) length in the indicated genotypes. Data are mean MV length/organoid (N=50 MV/organoid, 5-10 organoids). n.s. non-significant, *p<0.05, ****p<0.0001. (C) Quantification of the number of F-actin^+^ rounded foci on the full volume of an organoid, N= 27-49 organoids per condition from 3 independent experiments. (D-E) *Atp6v0d1* KO and *Myo5b* KO induce MVIs (D) visualized by F-actin staining (left) and TEM (right, arrows), as well as ectopic lumen (E, arrows). Inserts in A) and D) are magnified images showing the brush border and MVIs, respectively.

*MYO5B, STX3, STXBP2/MUNC18*.*2* and *UNC45A* encode factors implicated in the recycling of apical proteins through Rab11^+^ endosomes^1, 3^. Consistently, as observed by TEM and Periodic acid Schiff (PAS) staining, *Myo5b* KO organoids displayed a subapical accumulation of tubulo-vesicular compartments, and of the apical transmembrane proteins CD10 and DPPIV, or the trafficking factors Rab11 and STX3 (Figure 2A-C and Supplementary Figure 2A-F). Surprisingly, while V_0_-ATPase controls the same recycling step in *C. elegans*^4^, V_0_-ATPase KO in organoids was not associated with defective apical trafficking, suggesting that its function on apical membrane homeostasis differs from *bona fide* MVID factors (Figure 2A-C and Supplementary Figure 2A-F).

**Figure 2.**
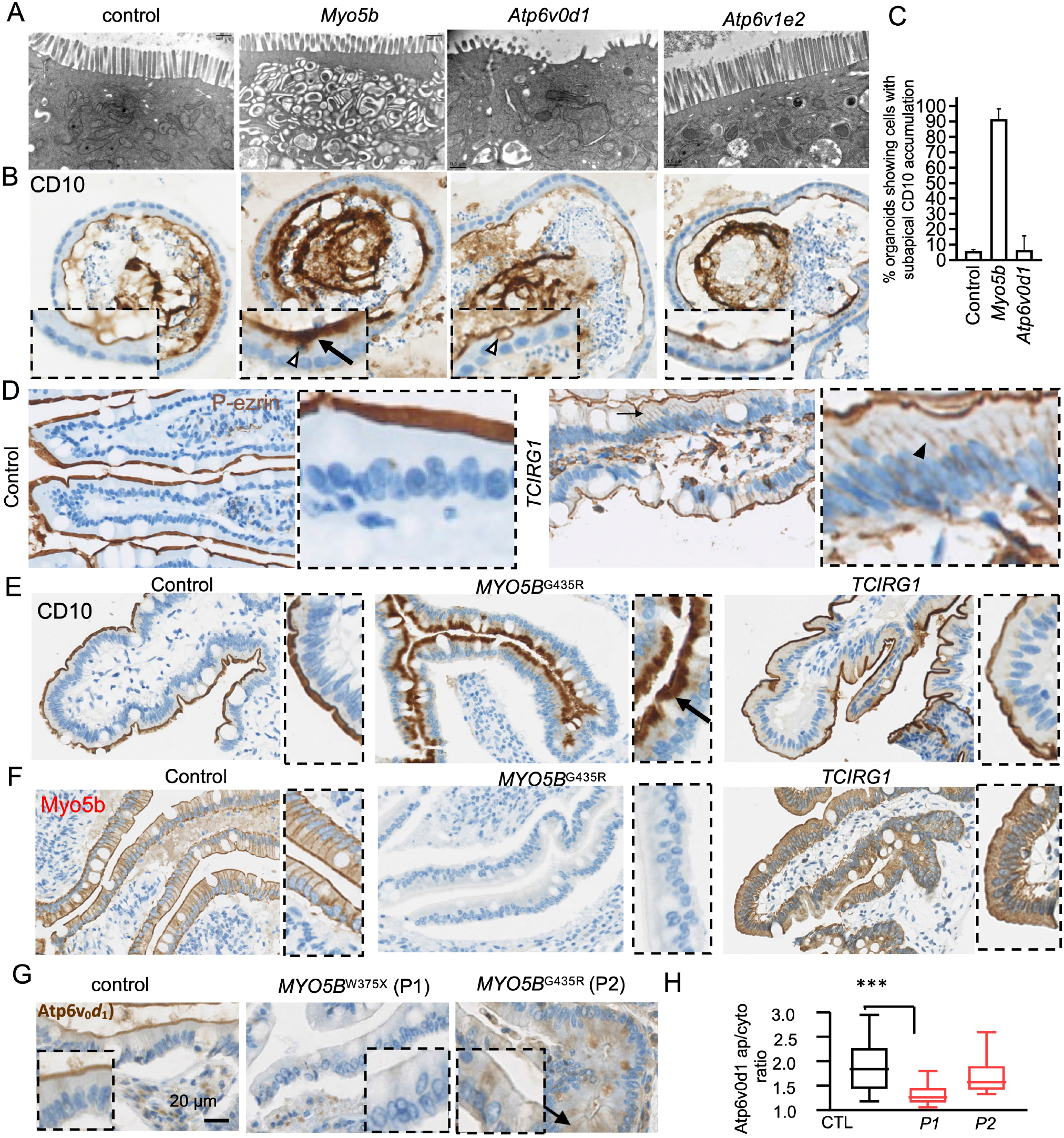
Comparison of Myo5b and V_0_-ATPase function on brush border, polarity and trafficking. (A) TEM analysis of the subapical cytoplasmic content in the indicated organoids. (B-G) IHC staining of phospho-ezrin (P-ezrin) (D), CDlO (B-C, E), Myo5b (F) and Atp6v0d1 (G) in mouse intestinal organoids (B) or human duodenum samples (D-G). (C) Quantification of the number of organoids displaying a subapical accumulation of CD10 (N=33-49 organoids from 2 independent experiments). (H) Quantification of Atp6v0d1 apical/cytoplasmic ratio (N=10 villi). Inserts are magnified images. Arrows, open and closed arrowheads indicate the subapical accumulation of markers, MVIs, and the basolateral appearance of markers, respectively. ***p<0.001, unpaired student t-test.

To confirm these results in humans, we analyzed small intestine resections from a patient suffering from osteopetrosis, a rare disease caused by mutations in the V_0_-ATPase *a*3 subunit-coding gene *TCIRG1*^9^ (Supplementary Figure 3A). IHC against the BB marker phospho-ezrin revealed its basolateral mislocalization in this *TCIRG1* patient compared to control (Figure 2D and Supplementary Figure 3B) similarly to its fate in MVID patients mutated for *MYO5B*^10^ and confirming the polarity defects observed in organoids (Figure 1E, Supplementary Figure 3C-D). However, this BB defect was not associated with a subapical accumulation of PAS nor apical transporters, unlike in a *MYO5B* patient, confirming our observations in organoids (Figure 2E and Supplementary Figure 3E-F). Thus, our data demonstrate that disruption of V_0_-ATPase function induces an atypical MVID phenotype where a microvillus atrophy is uncoupled from apical trafficking defects. Like many patients with infantile osteopetrosis, this TCIRG1 patient presented failure to thrive and an important feeding disorder requiring enteral nutrition through a gastrostomy for many years. However, there was no obvious intestinal absorption defect, indicating that a microvillus atrophy may not be sufficient to provoke nutrient absorption defects and suggesting that MVID symptoms are likely the consequence of both a BB atrophy and a failure in apical protein targeting.

Finally, to test the putative link between V_0_-ATPase and MVID, we studied the mutual requirement between Myo5b and Atp6v0d1 for their apical localization. Myo5b, which localized at the cell cortex in control samples, partly accumulated in the cytoplasm but remained apically localized upon *TCIRG1* mutations (Figure 2F). Contrarily, Atp6v0d1 apical localization was dramatically affected in two MVID patients carrying *MYO5B* mutations (Figure 2G-H), suggesting that the microvillus atrophy associated with MVID could be due to the loss of the V_0_-ATPase. We therefore propose that MVID might be induced by the combination of two independent processes: microvillus atrophy and apical transport defects.

## Abbreviations

MVID: microvillus inclusion disease
MVI: microvillus inclusion

## Acknowledgments

We thank Hans Clevers for the Noggin and R-Spondin1 expressing cells, Caroline Poix for preliminary imaging, Guillaume Halet for his help with organoid culture set up and members of the Michaux lab for discussions. IHC and imaging were performed at the Histo pathology High precision (H2P2) and the Microscopy Rennes Imaging Center (MRic Photonics and TEM) facilities of the UMS Biosit, member of the national infrastructure France-BioImaging supported by the French National Research Agency (ANR-10-INBS-04).

---

**Supplementary Figure 1.**
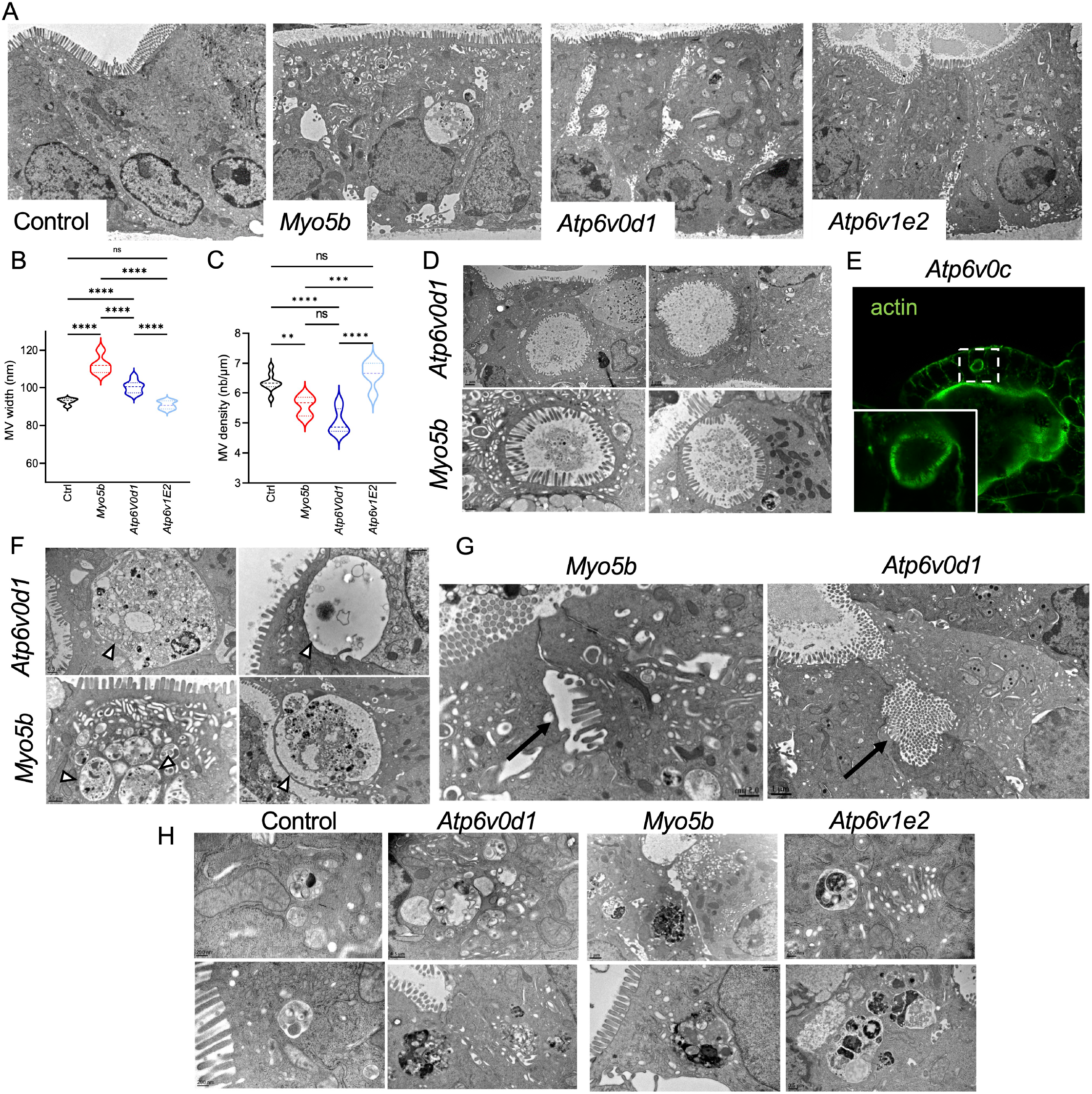
MVID-like structural defects. (A) Overview of the enterocytes observed by TEM in the indicated KO organoids. (B-C) Quantification of MV witdh (B, N=50 MV/organoid, 5-10 organoids) and density (C, N=10 MV/μm/organoid, 5-10 organoids) from TEM images. (D) Additional images of the MVIs induced by *Atp6v0d1* and *Myo5b* KO in organoids. (E) KO of the V_0_-ATPase subunit *Atp6v0c* also induces the appearance of F-actin^+^ MVIs. The insert is a magnified view of the selected area. (F-G). Knockout of both *Atp6v0d1* and *Myo5b* induce mixed organelles (F, arrowheads) and lateral microvilli (G, arrows). (H) Abnormal lysosomes are observed upon V_0_- *(Atp6v0d1)*, V_1_-ATPase *(Atp6v1e2)* and *Myo5b* KO.

**Supplementary Table 1.** Phenotypic comparison between control, *Atp6v0d1, Myo5b* and *Atp6v1e2* KO organoids.

| Sample | Control | *Atp6v0d1*-KO | *Myo5b*-KO | *Atp6v1e2*-KO |
| --- | --- | --- | --- | --- |
| BB/Microvilli | Normal | **Atrophy (100%)**<br>Variable atrophy of brush-border MV, local BB denudation, altered dimensions of apical MV | **Atrophy (100%)**<br>Variable dystrophy of brush-border MV, altered dimensions of apical MV | Normal |
| Subapical vesicular structures | No accumulation | No accumulation | **Subapical accumulation** of different vesicles and tubules ("secretory granules") (100%) | No accumulation |
| Enlarged vacuolar structures/Mixed organelles | No | **Abnormal vesicle-like (mixed) organelles (100%)**<br>various sizes, shapes, and electron density, with components of immature granules and lysosomes | **Abnormal vesicle-like (mixed) organelles (100%)**<br>various sizes, shapes, and electron density, with components of immature granules and lysosomes | No |
| Microvillus inclusions (MVIs) | No | **MVIs (40%)** lined by inward-pointing microvilli located in the apical/middle cytoplasm | **MVIs (100%)** lined by inward-pointing microvilli located in the apical/middle/basal cytoplasm | No |
| Lysosomes | Small and normal (33%) | **Giant and abnormal (100%)**<br>Non-functional autolysosomes with heterogeneous sizes and contents, Acidification/degradation defects | **Abnormal (100%)**<br>Non-functional autolysosomes with heterogeneous sizes and contents, Acidification/degradation defects | **Abnormal (100%)**<br>Non-functional autolysosomes with heterogeneous sizes and contents, Acidification/degradation defects |
| Ectopic lumen | No | **Lateral lumen with microvilli-like structures (50%)** | **Lateral lumen with microvilli-like structures (20%)** | No |
| Intercellular spaces/Lateral interdigitations | (22%) | **Widened lateral spaces with abundant lateral interdigitations, as well as bundles of ectopic microvilli facing laterally (100%)** | **Widened lateral spaces with abundant lateral interdigitations, as well as bundles of ectopic microvilli facing laterally (100%).** | **Widened lateral spaces with abundant lateral interdigitations, as well as bundles of ectopic microvilli facing laterally (100%).** |
| Basal membrane | Normal | **Basal interdigitations (100%)** | **Basal interdigitations (90%)** | **Basal interdigitations (90%)** |

**Supplementary Figure 2.**
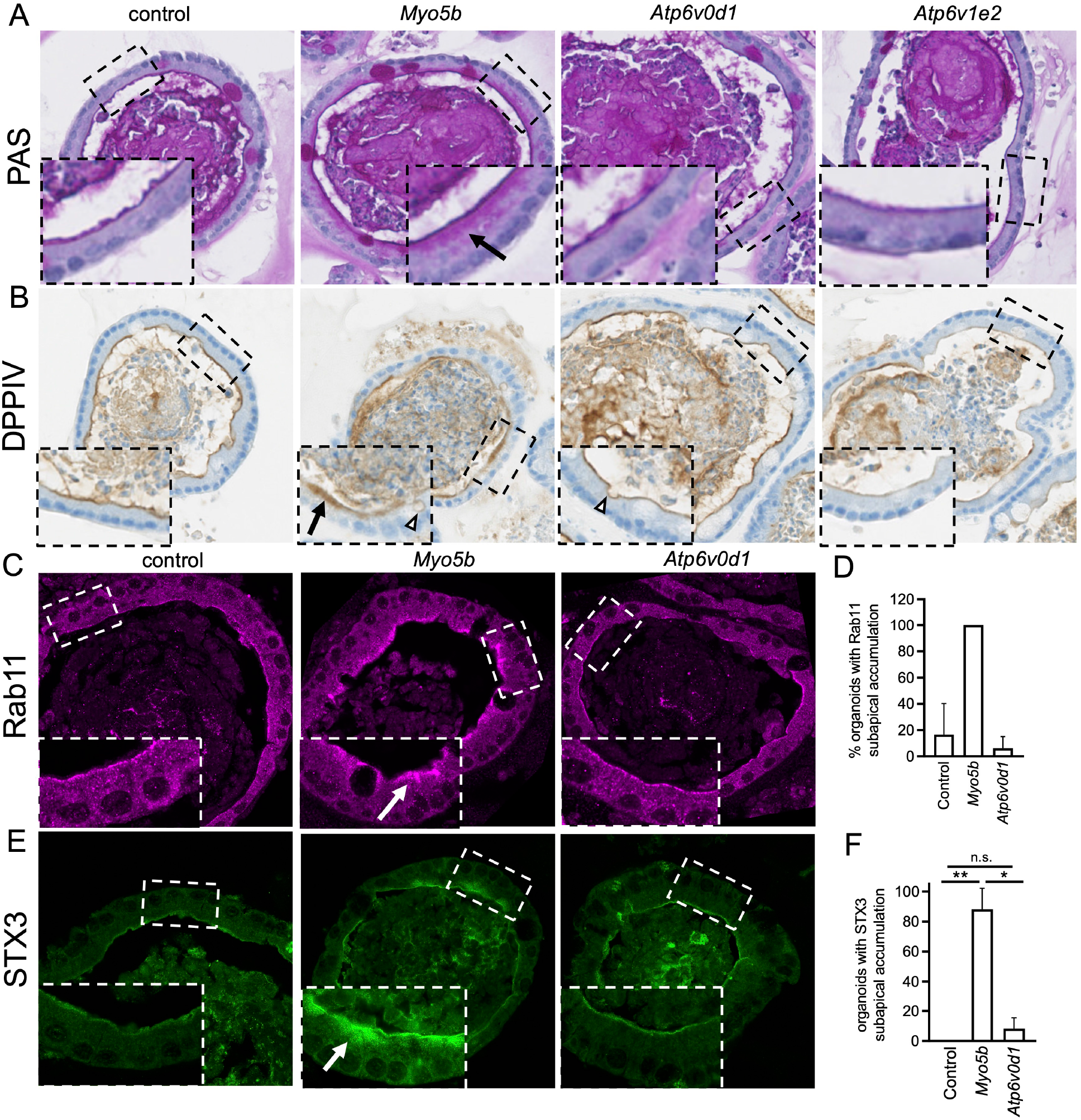
Analysis of apical membrane trafficking. Mouse intestinal organoids were stained with periodic acid Shiff (PAS) (A) or with antibodies directed against DPPIV (B), Rab11 (C) or STX3 (E). (D-F) Quantification of the number of organoids presenting cells with a subapical accumulation of Rab11 (D, N= 23-28 organoids from 2 independent expermients) and STX3 (F, N= 23-32 organoids from 3 independent experiments). Inserts are magnified images of the indicated ROIs. Arrows, open and closed arrowheads indicate the subapical accumulation of markers, MVIs, and the basolateral appearance of markers, respectively. n.s., non significant; *p<0.05, **p<0,01.

**Supplementary Figure 3.**
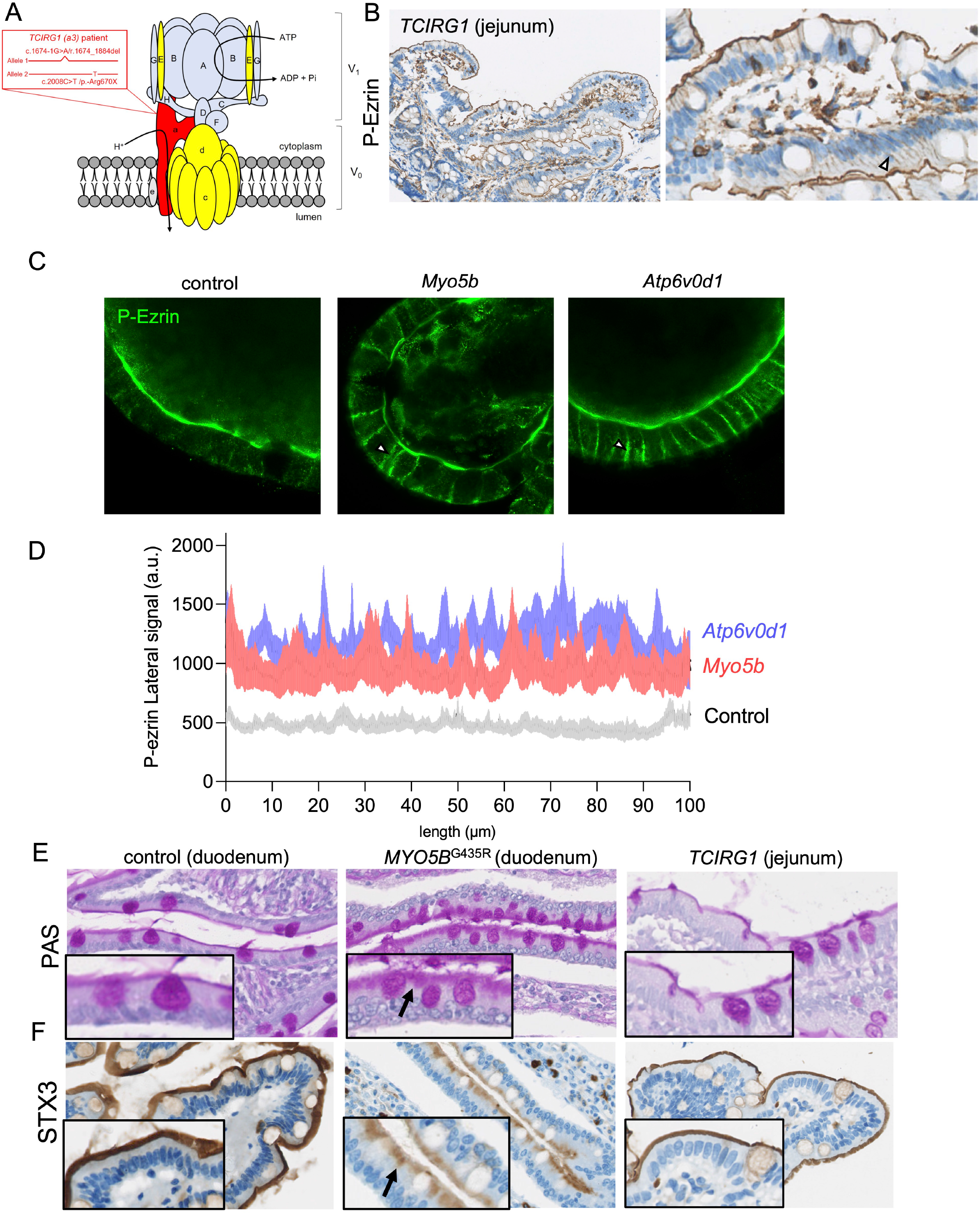
Conservation of V-ATPase function in human samples. (A) Compound heterozytgote mutations affect *TCIRG1* transcript’ s splicing (c.1674-lG>A) and translation (c.2008C>T). (B) IHC directed against Phospho-ezrin on jejunum samples from the patient carrying *TCIRG1* mutations. (C-D) The indicated organoids were stained for phospho-ezrin. (C) shows representative images and (D) the mean ± SEM of phospho-ezrin signal along a 100 μm line crossing the lateral membranes of cells (N= 10-14 organoids). (E-F) Intestinal samples from patients with MVID (MYO5B^G435R^) or carrying mutations on *TCIR11* were stained with PAS (E) or with antibodies directed STX3 (F). Inserts are magnified images. Arrows, and arrowheads indicate the subapical accumulation of markers and the basolateral appearance of markers, respectively.

